# Mothers facing greater environmental adversity experience increased costs of reproduction

**DOI:** 10.1101/2024.12.24.630228

**Authors:** Euan A. Young, Erik Postma, Virpi Lummaa, Hannah L. Dugdale

**Author notes:** equal last authorship (Virpi Lummaa, Hannah L. Dugdale).

## Abstract

Evolutionary theory of aging predicts that women with increased reproductive effort live shorter lives, but evidence is inconsistent. These inconsistencies could be because environmental conditions influence how much a mother’s life span is reduced when having more children, i.e. their life-span cost of reproduction. Using a structural equation measurement model, we compare how reproductive effort affects the life span of 4,684 women exposed across different life-stages, or not at all, to the Great Finnish Famine. We find that life span costs of reproduction became higher in mothers exposed to the famine during reproduction, and for these mothers amounted to lower life expectancy of ∼0.5 years per child. Conversely, reproduction did not shape the life spans of mothers not exposed to the famine, or exposed post-reproduction or during development. This natural experiment reveals how environmental adversity can influence reproductive costs, providing a biological explanation for previous inconsistent findings, while showing how reproductive behavior has shaped the evolution of aging in humans.

**TEASER:** Mothers with more children had shorter lives during a famine, suggesting that harsh conditions increase costs of reproduction.

## INTRODUCTION

Understanding why we age – defined as a physiological decline in later-life – and why some of us undergo this aging process faster than others, are vital foundations for improving human health-spans (1). One theory – the disposable soma hypothesis – predicts that aging is a consequence of natural selection honing our physiology to maximize reproduction at the cost of other organismal functions, resulting in an accumulation of damage through life that limits lifespans (2,3). Consequently, researchers have tested if individuals who devote more resources to reproduction have shorter lifespans (4). However, expected negative associations between reproduction and lifespans are rarely found (5,6) and typically only when using experimental manipulations (e.g., (7)). For humans, the bearing and raising of even one child is energetically demanding for women (8), but – despite over 100 years of research (9) – evidence of a reproduction-survival trade-off is inconsistent (10–12) and the role that human reproductive behavior plays in shaping aging remains unclear.

One potential explanation for these inconsistencies is that the reproduction-survival trade-off does not manifest itself in the same way when measured across a heterogeneous set of individuals (13). First, individual heterogeneity will weaken the observed slope of the regression of survival against reproduction (i.e. the estimate of the reproduction-survival trade-off *function*) because among-individual variation in resource acquisition introduces a positive association between reproductive effort and lifespan that can mask the within-individual reproduction-survival trade-off (i.e., the big house and big car syndrome (14,15)). Thus, the *estimated* trade-off function quantifies how important reproductive behavior versus individual heterogeneity is in shaping human lifespans, and studies aim to account for such individual heterogeneity in their analyses (16). Second, among-individual variation (e.g., in frailty or quality), due to intrinsic or environmental differences, may mean that some women will manifest costs of reproduction more than others (12,13,17–19). Previously, the reproduction-survival trade-off has been found to be stronger in lower socioeconomic status women (20,21) and women experiencing higher levels of infant mortality (22). However, we still have a poor understanding of which biological factors drive these changes in the reproduction-survival trade-off, with studies showing considerable variation in the estimated trade-off within the same populations over time, in line with environmental causes playing a role (23–25). Understanding these drivers can give us insight into the mechanisms underlying this fundamental trade-off, how it manifests itself, and under what circumstance human reproductive behavior is an important factor in shaping aging.

Although the role of reproductive effort in shaping aging is uncertain, examples of environmental effects on human health are numerous (26,27). In humans, experiencing periods of extreme environmental adversity, such as famines, can accelerate the biological aging process (28). In fact, famine can have an array of effects, from long-term individual effects on human health, such as prenatal famine exposure increasing risk of type-2 diabetes and schizophrenia later in life (29,30), to social and economic consequences (31). Although widely used as natural experiments in other fields, the role that famine exposure plays in shaping estimated lifespan costs of reproduction has not to our knowledge been studied.

Given the well-documented adverse effects of famines, mothers exposed to famines could show greater reductions in lifespan for a given increase in reproductive effort. However, this reduction may depend upon the life-history stage of the mother during the famine exposure. Firstly, even short-term fluctuations in early-life environmental conditions can lead to physiological differences (32) that influence the overall lifespan and realized reproduction of individuals (33,34). Thus, women experiencing famines during development may show increased costs of reproduction. However, Nenko et al. (24) found no evidence that crop yields, spring temperatures, or infant mortality rates at birth modified the reproduction-lifespan trade-off in mothers. Instead, mothers may be especially vulnerable to resource limitations while raising children (24), due to both the calorific demands of pregnancy and lactation, and increased vulnerability to disease (8). Indeed, Wang et al. (25) proposed that some individuals experiencing the Great Depression and Second World War while raising and bearing offspring caused the variation in costs of reproduction in the Framingham cohort study. Finally, women in post-reproductive life are particularly at risk of mortality under lower food availability (34). Thus, mothers could suffer increased frailty due to high reproductive effort, but perhaps only when combined with famine exposure during post-reproductive life would this result in reduced lifespans. However, no study has examined how a period of environmental adversity affects lifespan costs of reproduction in humans and whether this might vary across life-stages.

Rigorous tests of these hypotheses require detailed individual-level life-history data for a period encapsulating a major famine event. Here, we use individual life-history data across 250 years from rural Finland to quantify lifespan costs of reproduction and compare how they changed across mothers exposed to the Great Finnish Famine in different life-stages, and those not exposed at all. In the 1860s, Finland experienced several harsh winters resulting in a series of low harvests, which in 1867 dropped to catastrophic levels, resulting in a major loss of employment (35). Widespread migration then followed as people went in search of work, which in turn triggered a large spike in infectious diseases, such as typhoid fever, typhus, and dysentery (35). This led to an excess mortality of 110,000 (or 5-10% of Finland’s population (36)) making it one of the most devastating famines in recent European history, and the worst in Finland since the 17^th^ century (37).

Here we capitalize on the fact that this famine occurred in the middle of our study period, allowing us to estimate the lifespan costs of reproduction before, during, and after a famine. Specifically, we compare mothers exposed to the famine during three life-stages (during their development [ages 0-19], while reproductively active [ages 19-45], and while in post-reproductive life [ages 45+]) to mothers dying before the famine and those born in the years after using a structural equation measurement model (N = 4,868, Fig. 1). We follow Helle (38) and use a measurement model using multiple indicator variables to obtain a comprehensive measure of lifetime reproductive effort. Within this structural equation model framework, we estimate the association between reproductive effort and lifespan (i.e., the reproduction-survival trade-off function) in each famine exposure group and how these lifespan costs of reproduction differed across groups, accounting for the changes in reproductive effort and lifespan in relation to famine exposure. Broadly we expected mothers exposed to the famine to show an increased lifespan cost of reproduction, but that this cost would depend upon the life-stage of mothers during this famine exposure. Showing such a dependency would shed light on why the estimated costs of reproduction are so variable both among and within studies.

**Fig. 1:**
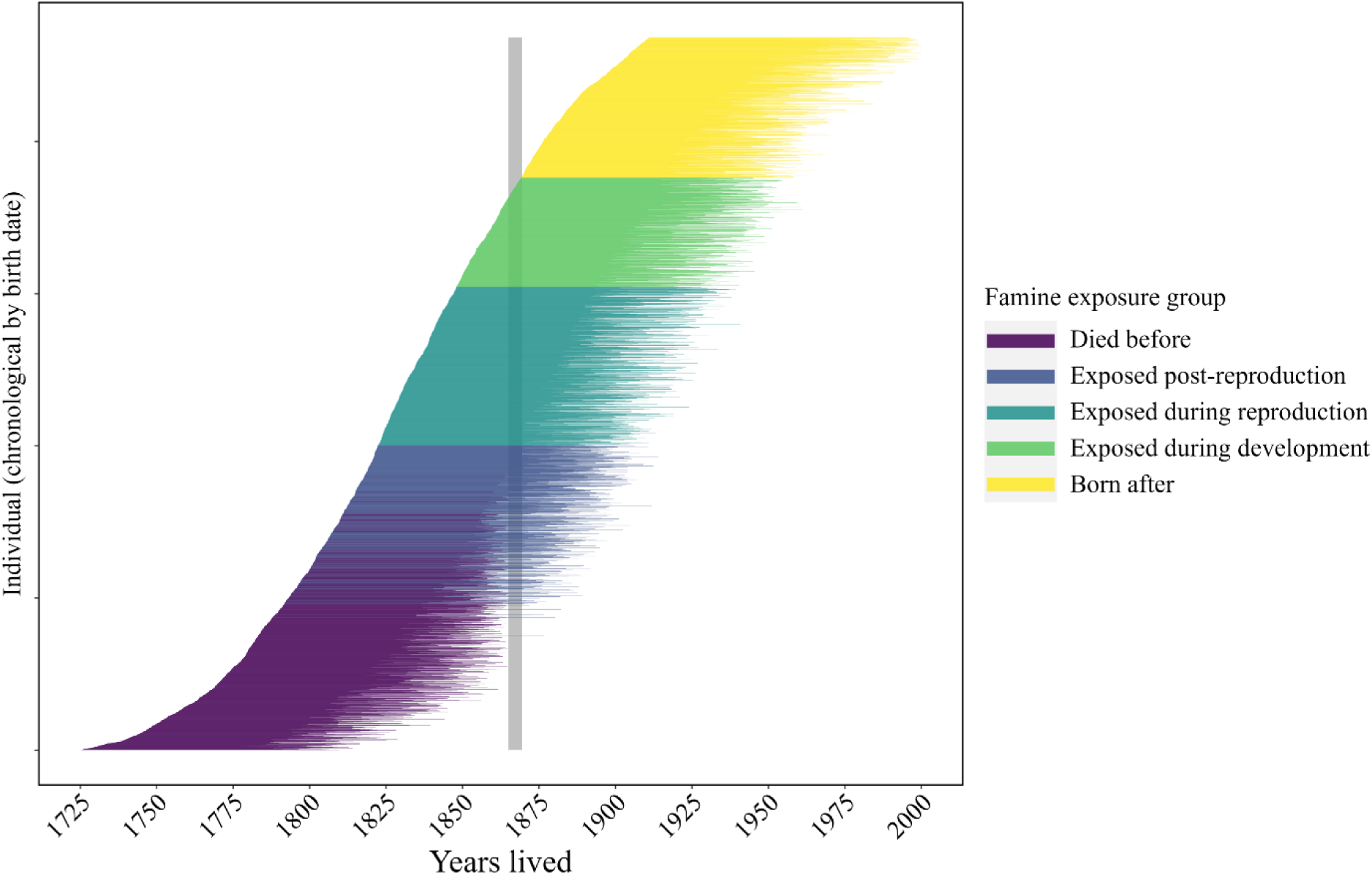
Line graph showing the years lived for each mother included in the analyses, sorted chronologically by their birth date (N = 4,684). Colors indicate their exposure to the Great Finnish Famine (grey bar) and groupings in the analyses: died before (n = 1,297), exposed post-reproduction (n = 705), exposed during reproduction (n = 1,044), exposed during development (n = 716), and born after (n = 922).

## RESULTS

### Lifespan costs of reproduction

Exposure to the Great Finnish Famine shaped the estimated reproduction-survival trade-off (Fig. 2, Tables 1 and 2). Specifically, increases in reproductive effort were associated with a reduction in lifespan in mothers exposed to the famine while reproductively active, but not among other groups (Fig. 2; Table 2). Mothers exposed to the famine who had one child would be expected to live to age 71.6 (95% CIs 69.2 – 74.6), whereas a mother with 15 children would only be expected to live to age 64.3 (61.7-68.3), or a reduction in lifespan of half a year per child (Fig. 2, Table S1). In line with this, pairwise comparisons of the five groups showed that the association between reproductive effort and lifespan in mothers exposed during reproduction was also higher than in mothers dying before the famine, mothers born after the famine, and mothers exposed to the famine in post-reproductive life (p < 0.05), and marginally higher than those exposed to the famine during development (p = 0.087, Table 2, Fig. 2).

**Fig. 2:**
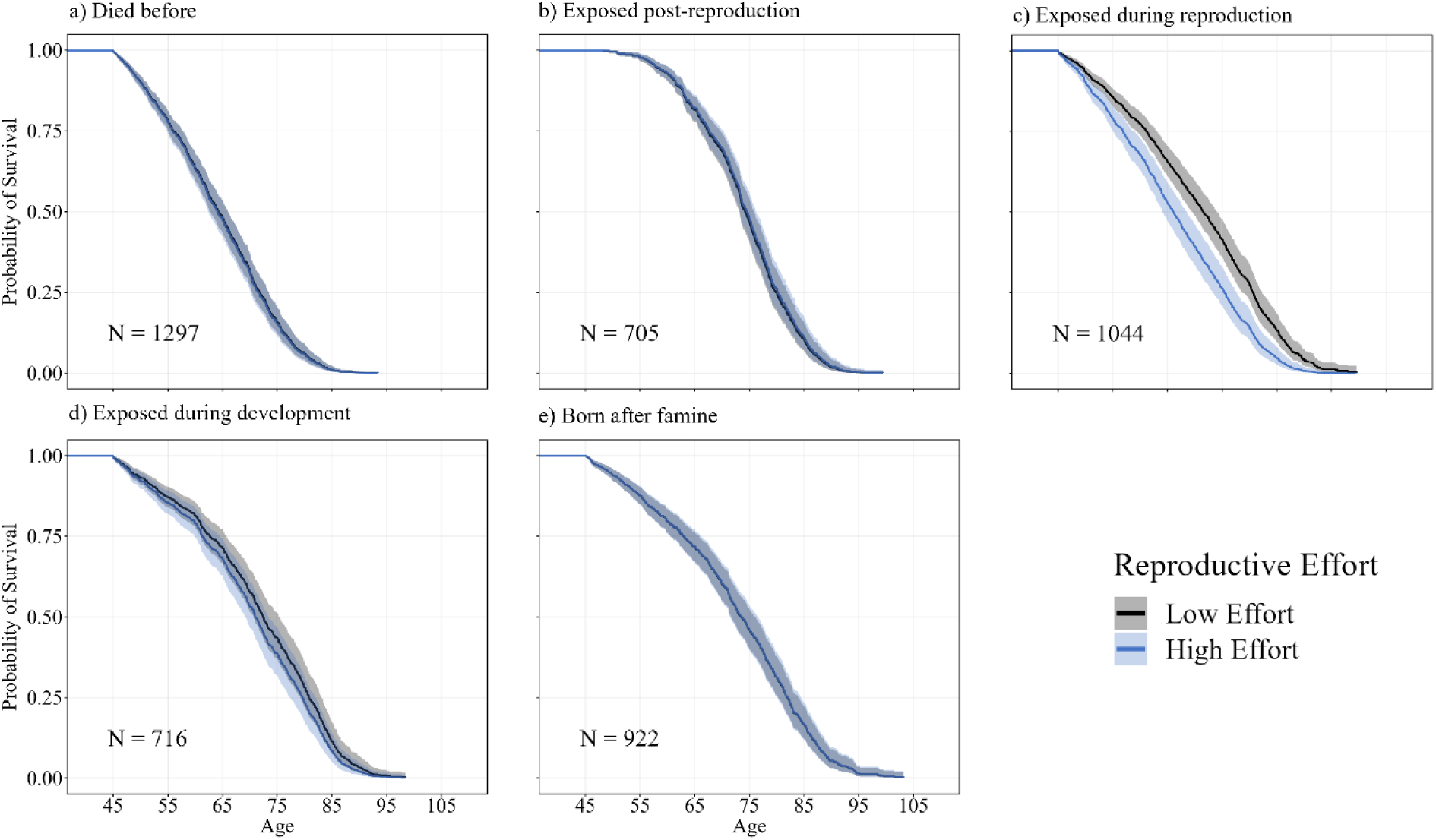
Predicted survival curves for low and high reproductive effort across five famine exposure groups. Curves were generated using Cox proportional hazards models on each famine exposure group separately (see Table S1). Each panel represents a different famine exposure group: (a) died before famine, (b) exposed during post-reproduction, (c) exposed during reproduction, (d) exposed during development, and (e) born after famine. Survival curves are shown for low (5th percentile, -3.8) and high (95th percentile, 6.5) reproductive effort levels. All curves are calculated for individuals of intermediate socioeconomic status and median relative birth date within each famine group. Shaded areas represent 95% confidence intervals.

**Table 1:**
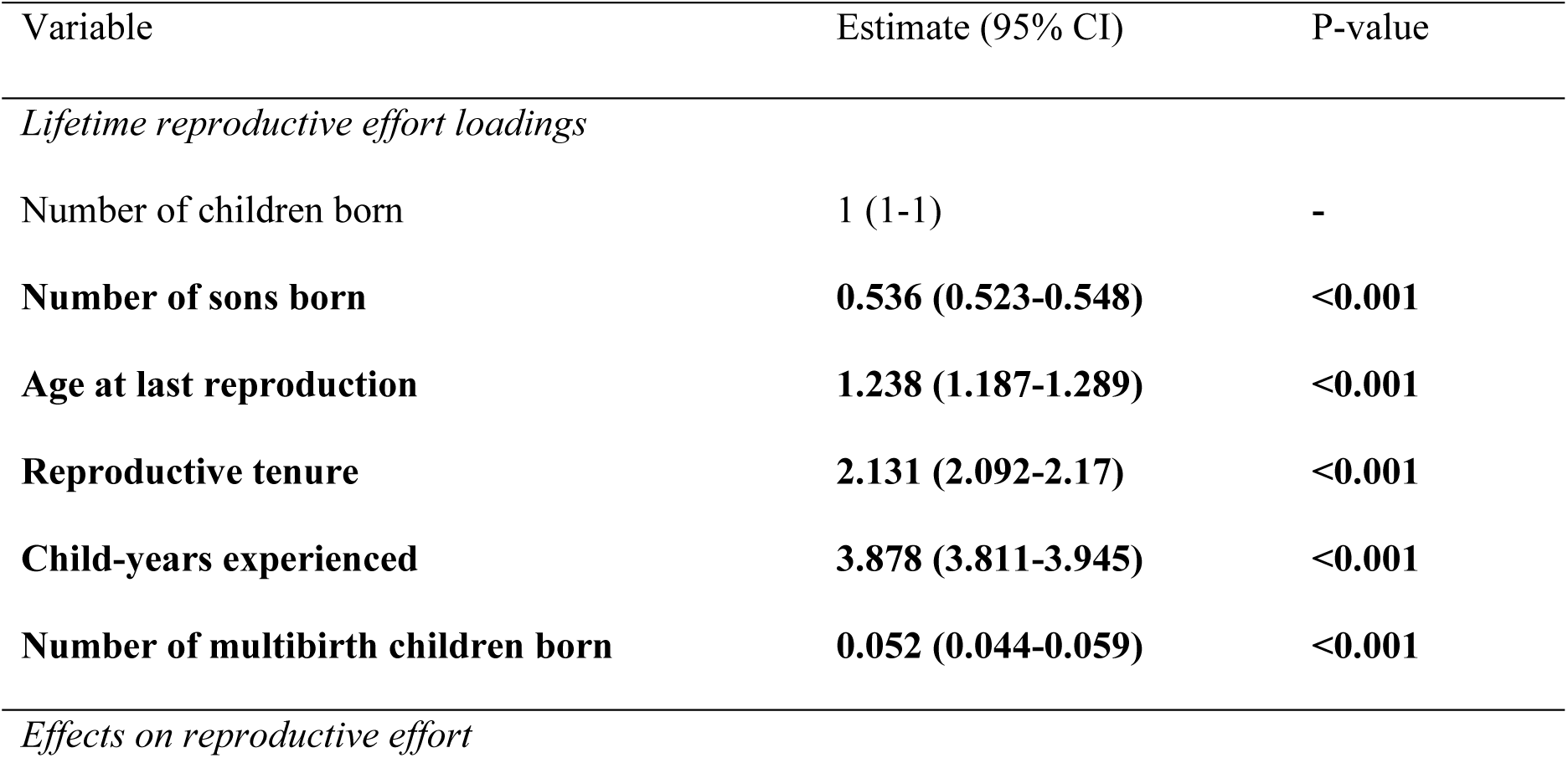
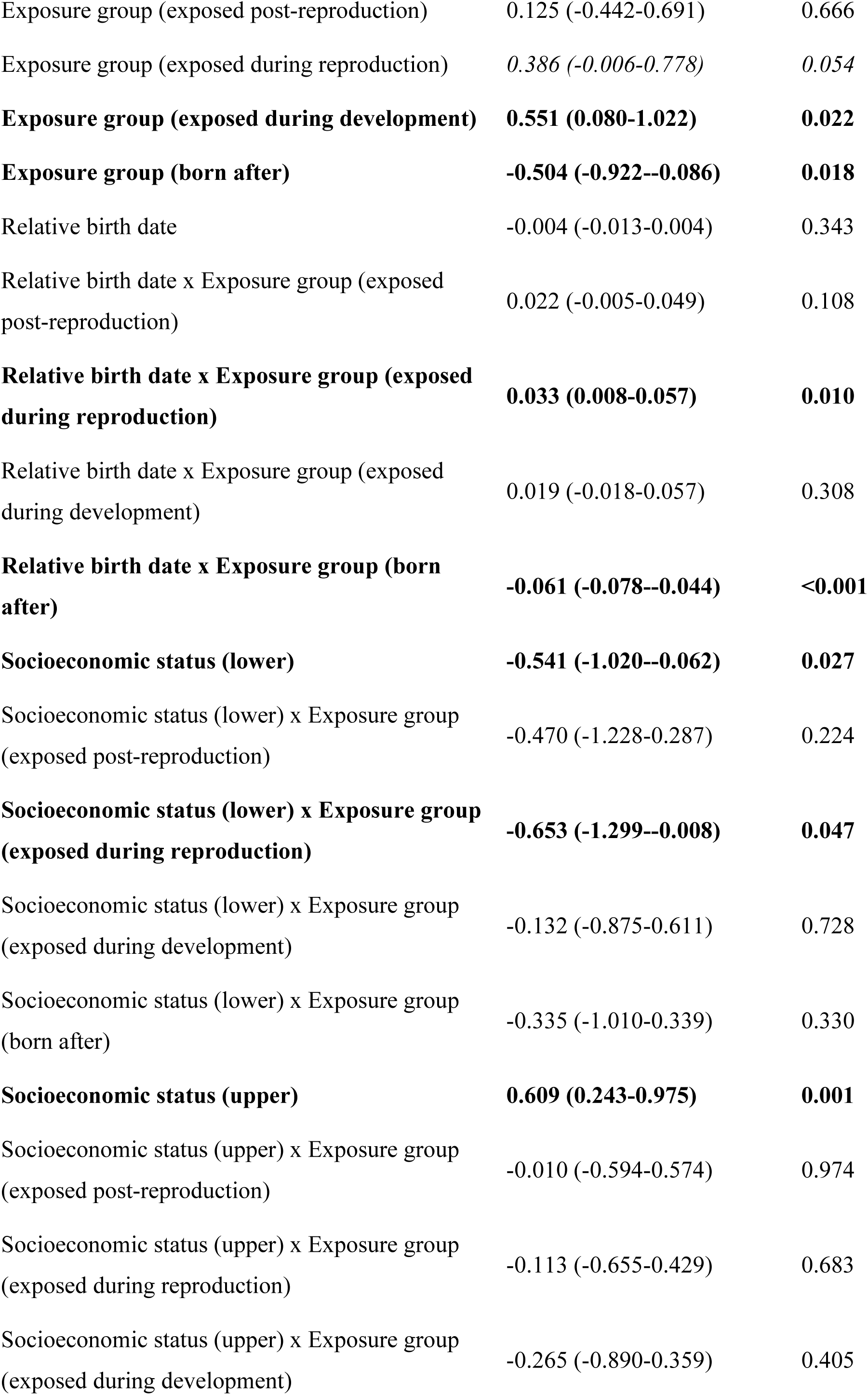
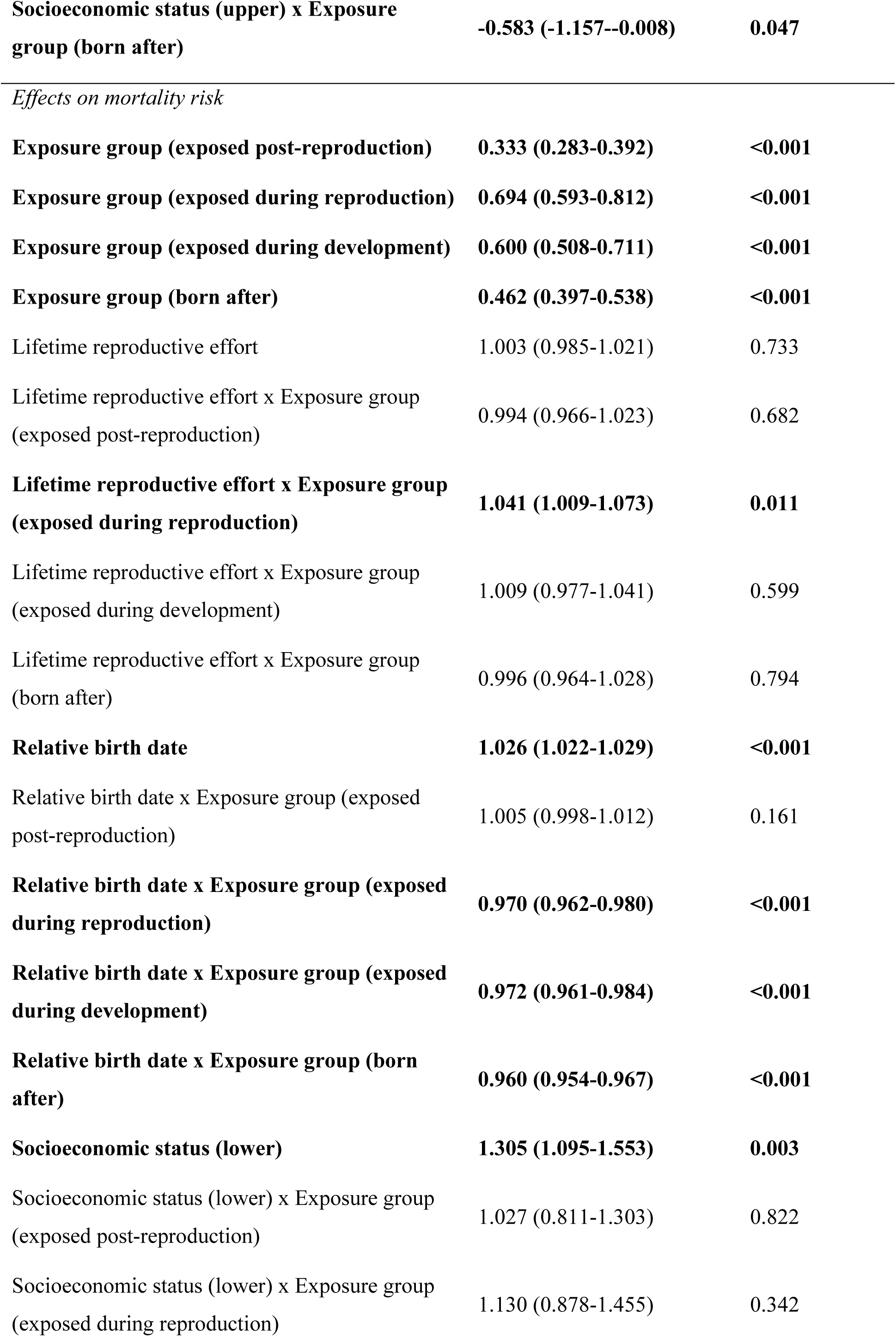
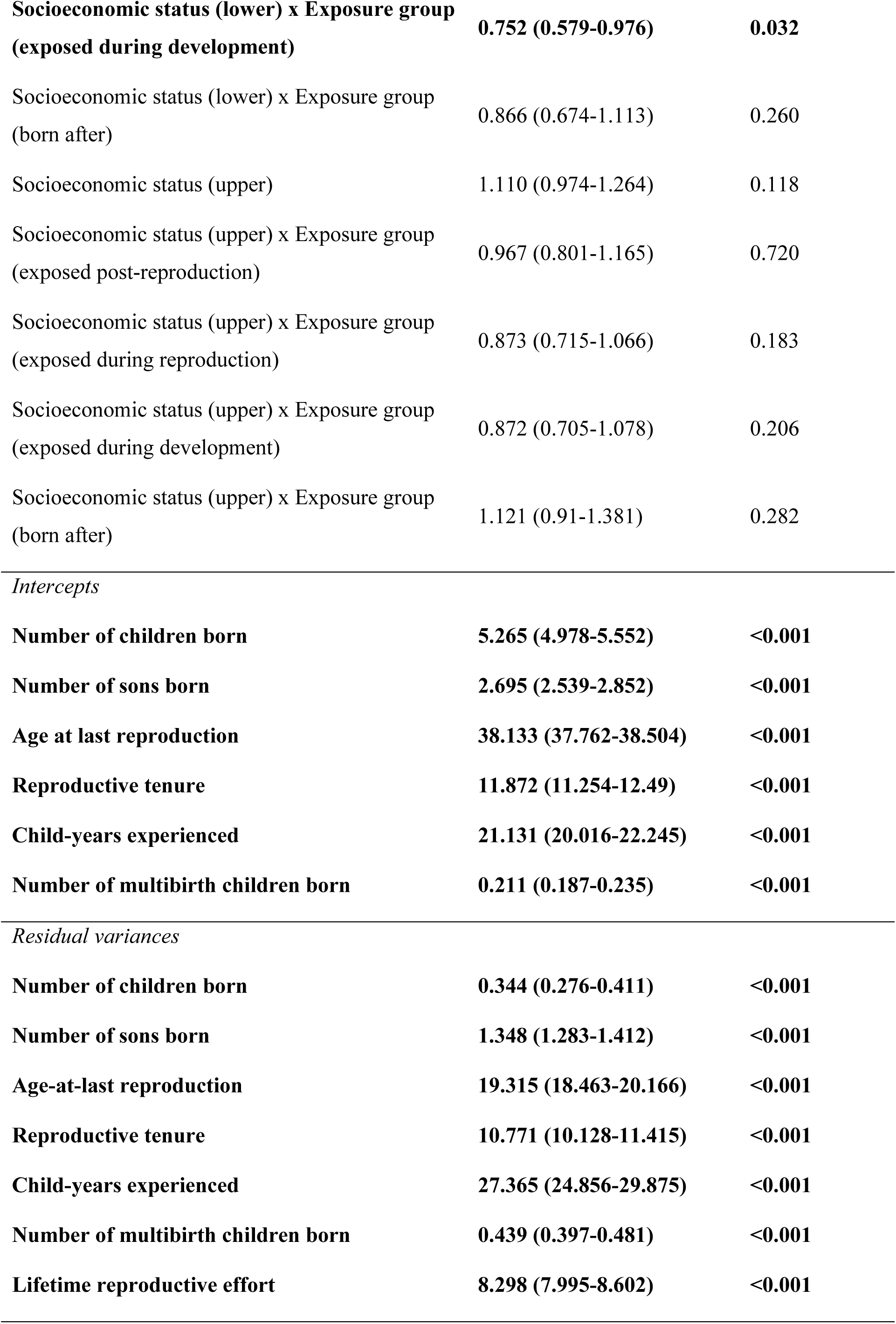
Structural equation model showing effects on lifetime reproductive effort and lifespan (as measured by log-hazard mortality risk) from 4,684 mothers. Factor loadings, mean values (*Intercept*), and residual variance for indicator variables used to estimate lifetime reproductive effort in the measurement model are shown. Estimates are shown with 95% confidence intervals and p-values. Significant effects (p < 0.05) are in bold. Effects of famine exposure are compared to women who died before the famine. Socioeconomic statuses are compared to intermediate statuses. Relative birth date is scaled by year and centered by the middle birth date for each group.

**Table 2:** Pairwise comparisons of the association between reproductive effort and mortality risk across famine exposure groups. Diagonal values show the hazard ratio (95% Confidence Intervals) for the effect of reproductive effort on mortality within each group. Off-diagonal values show p-values for pairwise differences in this association between groups. Significant differences or associations (p < 0.05) are highlighted in bold; marginally significant associations (p > 0.05 and < 0.10) are italicized. Results are from a structural equation model of the same structure as Table 1 but with different reference groups to estimate p-values and associations.

|  | Died before<br>famine | Exposed post-reproduction | Exposed<br>during<br>reproduction | Exposed<br>during<br>development | Born after<br>famine |
| --- | --- | --- | --- | --- | --- |
| **Died before**<br>**famine** | 1.003 (0.985<br>- 1.021) |  |  |  |  |
| **Exposed post-**<br>**reproduction** | 0.682 | 0.997 (0.975 -<br>1.019) |  |  |  |
| **Exposed**<br>**during**<br>**reproduction** | **0.011** | **0.007** | **1.044 (1.018 -**<br>**1.069)** |  |  |
| **Exposed**<br>**during**<br>**development** | 0.599 | 0.404 | *0.087* | 1.012 (0.986 -<br>1.039) |  |
| **Born after**<br>**famine** | 0.794 | 0.926 | **0.018** | 0.5 | 0.999 (0.972 -<br>1.026) |

These results were confirmed in a complementary mixed model analysis that accounted for significant variation in lifespan amongst birth years, families, and 13 geographical regions (Table S2). Mothers again varied in the reproduction-survival trade-off function according to famine exposure, with the statistically significant association between reproductive effort and lifespan only found in mothers exposed to the famine during reproduction, and similar in magnitude as estimated in the previous model (Tables S3 and S4). The association between reproductive effort and lifespan in mothers exposed to the famine while reproductively active was also higher than in all other groups, except for those exposed during development (Table S4).

### Temporal changes in lifespan

There were differences *among* famine exposure groups in mean lifespan, and in how lifespan changed over time *within* each group (Table 1, Table S1). Life-expectancy for mothers dying before the famine was 66 years for those born in 1777 but declined over time to 55 years for mothers born in 1819 (Table 1, Table S1). Similarly, for mothers exposed to the famine post-reproduction, life-expectancy was 79 years for those born in 1796 but declined overtime to 70 years for mothers born in 1822 (Table 1, Table S1). These trends reflect the classification of these groups: within mothers who died before the start of the famine, those born later must have lived shorter lives. Similarly, by focusing only on famine survivors, mothers born earlier among those exposed to the famine during post-reproduction cannot have had short lives, with the higher life-expectancies overall likely reflecting the loss of higher frailty women during the famine itself. Mothers exposed to the famine during reproduction had a life expectancy of 71 years, which stayed similar for mothers exposed during development and did not change through time in either group (Table 1, Table S1). However, life-expectancies then increased for mothers born after the famine from 71 years in mothers born in 1869 to 78 years for mothers born in 1910 (Table 1, Table S1).

### Temporal changes in reproductive effort

Mean reproductive effort did not differ between mothers dying before the famine and those exposed to the famine post-reproduction, and it remained constant overtime within each group (Table 1, Fig. S1). However, for mothers exposed during reproduction, reproductive effort increased from being similar to prefamine levels for mothers born in 1822 to being 0.8 children higher in mothers born in 1848 in this group (Table 1). Reproductive effort then remained similarly high for mothers exposed during development who had on average 0.6 children more than pre-famine levels, which did not change over time (Table 1). Reproductive effort then declined rapidly in mothers born after the famine going from mothers born in 1869 having 0.8 children more than mothers dying before the famine, to mothers born in 1911 having 1.8 children fewer than prefamine levels (Table 1).

### Socioeconomic status effects

Socioeconomic status did not modify the association between reproductive effort and lifespan (Table S5). However, there were differences in both lifespan and reproductive effort across socioeconomic status groups, with the magnitude of these differences changing according to famine exposure (Table 1, Table S1). No differences in lifespans were found in mothers dying before the famine between higher and middle socioeconomic statuses, and these effects did not change across famine exposure groups (Table 1). However, lower socioeconomic status mothers had lower life-expectancies of around three years compared with intermediate socioeconomic status mothers (Table 1). This difference in life-expectancies between lower and intermediate socioeconomic status mothers remained similar across famine exposure groups, except in lower socioeconomic status mothers exposed to the famine during development who did not have lower lifespans than mothers of intermediate socioeconomic status (Table 1).

Upper socioeconomic status mothers dying before the famine had higher reproductive effort than mothers of intermediate socioeconomic status, amounting to 0.6 more children (Table 1). These differences gradually decreased through time until mothers born after the famine, where upper and intermediate socioeconomic status mothers had identical reproductive effort (Table 1). Mothers from lower socioeconomic statuses had lower reproductive effort amounting to 0.6 fewer children in mothers dying before the famine than those from intermediate socioeconomic status. This difference in reproductive effort between lower and intermediate socioeconomic status mothers remained stable, except for mothers exposed during reproduction, where the difference increased to 1.2 fewer children.

## DISCUSSION

Using life-history records for 4,684 Finnish women, we find evidence supporting adverse environmental conditions increasing the importance of reproductive behavior in shaping a mother’s lifespan (Fig. 2, Table 2). Specifically, mothers exposed to the Great Finnish Famine (1866-68) while reproductively active showed reduced lifespans of approximately 0.5 years per child as estimated through a structural equation measurement model. On the other hand, mothers not exposed to the famine (i.e., mother who died before, or were born after) or those exposed to the famine in post-reproductive life or during development did not pay a lifespan cost of reproduction. Although the effects of famines on human health and aging are well documented (28), with some evidence for environmentally-dependent costs of reproduction in animals (13,19), this is, to the best of our knowledge, the first evidence of a famine event modifying the reproduction–survival trade-off function in humans. These findings therefore add a biological explanation – to previously highlighted methodological explanations (detailed in: Helle et al. (39); Helle (40)) – for why several studies have shown variation in costs of reproduction across cohorts (23–25): fluctuations in environmental conditions experienced by mothers while reproductively active impact the trade-off function. This strengthens the evidence for the existence of lifespan costs of reproduction and gives insight into under what circumstances human reproductive behavior is an important driver of aging.

These results align with other studies showing variation in the estimated reproduction-survival trade off amongst mothers in human populations. For instance, one recent study using data for preindustrial Canada showed that only mothers having both high reproductive effort (i.e., a high number of sons) and high lifetime infant mortality showed increased lifespan costs of reproduction (22). While this change in the reproduction-survival function across mothers is in line with our findings, Invernizzi et al. (22) did not explore how environmental conditions drive variation in infant mortality among mothers beyond showing an effect of a mother’s birth year (see (22), Supplementary Table 1). These results also show parallels with studies showing that women of lower socioeconomic status showed a stronger trade-off (20,21). However, while these results align with some previous findings, we present the first evidence suggesting that environmental adversity influences the reproduction-survival trade-off across women.

Our finding that only mothers exposed to the famine while reproductively active showed lifespan costs of reproduction could be due to the increased energetic demands of bearing and raising children during a period of high environmental adversity, as previously suggested (41). When pregnant, mothers must increase their caloric consumption by on average 300 kcal/d to sustain a healthy pregnancy, which rises to 460-630 kcal/d during breastfeeding (42). If the mother’s pregnancy or breastfeeding coincides with a famine it could mean that these increases in metabolic demands are paid by themselves through lowering basal metabolism, and thus slowing or shutting down other important functions, such as immunity (11,43), resulting in a decline in health and shorter lifespans. Specifically, circulating estrogen levels – which are important for cardiovascular health (44) – are very low during breastfeeding (45). These levels are even lower in mothers with poor nutrition (46), which could mean that cardiovascular disease risk (which is already a risk factor of high parity (47)) could be even higher in mothers of high reproductive effort experiencing poor nutrition while reproducing. Although we can speculate, ultimately the underlying mechanisms are difficult to disentangle in genealogical data, but we hope that these results may help guide future human biobank studies or experiments on model species where unravelling underlying mechanisms is more attainable.

In addition to famine exposure, we found that demographic and cultural factors were important drivers of reproductive effort, lifespan, and their association. Differences across socioeconomic groups were pronounced, with lower socioeconomic status mothers having lower realized reproduction (37) and shorter lives (48), but not increased lifespan costs of reproduction (Supplementary Table S5, Helle (38), but see (21)). We also found that mothers exposed to the famine during reproduction lived shorter lives than those exposed during development, and those exposed during development lived shorter lives than mothers born after the famine. Rather than being a direct effect of famine exposure, this is probably a symptom of increased industrialization, which, in line with country-wide trends, started during this period (49). These improvements in lifespan through, for example, vaccination (50), combined with a reduction in overall reproductive effort, are potential additional reasons why the importance of the reproductive-survival trade-off decreased in mothers born after the famine. This is analogous to results from the Netherlands, where the reductions in lifespan for mothers having more children found in 1850 had disappeared by 1910 (51). However, these results require replication outwith northern European countries.

Although high reproductive effort during the famine may have affected individual health directly, among-individual processes, such as selective disappearance, may still have influenced the reproduction–survival trade-off function. Because the association between reproduction and survival will weaken with higher individual heterogeneity (14,15,17), higher mortality rates during the Great Finnish Famine may have driven higher mortality of frailer individuals (as suggested by a previous study (52)), reducing the overall amount of individual heterogeneity in mothers sampled after completing reproduction, and increasing the strength of the association. However, because our study necessarily included only survivors of the famine, selective disappearance due to the famine could have occurred in all of the famine exposure groups. Moreover, given that the effects of the famine on mortality were biased towards the very young and old, and lowest amongst reproductive age individuals (53), the selective disappearance should have been highest in the development and post-reproductive famine exposure groups – even after accounting for adverse conditions potentially increasing maternal death during childbirth (54). However, we found that the trade-off function was strongest and significant only in mothers reproducing during the famine, further suggesting that our results reflect a direct negative within-individual effect of famine exposure on reproducing mothers. Thus, while it is possible that among individual factors, such as selective disappearance, may have shaped some of our findings, the evidence suggests that it is not the primary cause of the stronger reproduction-survival trade-off in mothers exposed to the famine during reproduction.

However, higher selective disappearance in the mothers exposed during development may help explain why no moderating effects of the early-life environment have been found here and elsewhere (24). For instance, if selective disappearance removes the frailest mothers who are most susceptible to the trade-off, leaving a more robust subset of mothers for whom reproduction is less costly, the trade-off function may become weaker with greater selective disappearance. Some evidence of increased selective disappearance in the mothers exposed during development may be visible in our results, with women of lower socioeconomic status in general having lower lifespan, except if they have been exposed to the famine during development where selective disappearance may be larger. While evidence for cohorts in gestation during the Great Finnish Famine having shorter lives is lacking (55), given the importance of the early-life environment on human health (56), and in modifying the reproduction–survival trade-off function in wild animal populations (57), future studies should aim to better account for selective disappearance when studying how environmental adversity may modify lifespan costs of reproduction, in particular when looking for early-life effects.

We emphasize that our findings should be interpreted within the context of this being an observational study occurring over a single famine event that will have coincided with other historical events that may have shaped reproduction and lifespan. Albeit the largest (as measured by death rates (37)), the Great Finnish Famine is only one of many societal perturbations faced by Finns between the 18^th^ century and present day. Principally, the Finnish War (1808-1809), severe epidemics and crop failures (1832–1833), the Finnish Civil War (1918), and World War II may all have shaped the reproduction and longevity of individuals in our study population (58). Although we were able to account for some demographic trends in reproductive effort and lifespan both within and among famine groups, including expanding analyses to account for correlations among birth years, regions, and families, which did not affect results, there are limitations. Most importantly, because Finland began to industrialize soon after the famine (i.e. in the 1870s), we had no group of women born after the famine whose demography was comparable to the time preceding the start of industrialization. Thus, proving the causality of the Great Finnish Famine (1866-68) shaping the trade-off function is difficult within our system, but it may be possible in other datasets.

After over 100 years of research, human observational studies have provided mixed evidence for lifespan costs of reproduction, leading some to believe that reproductive behavior is not important in shaping human aging. Contrary to this, we provide evidence for a life-history trade-off between reproductive effort and lifespan and demonstrate that – under harsh conditions – reproductive effort is an important driver of individual variation in lifespan. These findings suggest that famines can provide fitness costs to mothers in the form of reduced lifespan, but may also include reduced reproduction of the children born from pregnancies during famine, although whether early-life environmental effects cause positive or negative effects fitness effects in humans remains debated (59,60). Although reproductive behavior may shape the lifespans of mothers exposed to adverse environments while reproducing, it appears less important in contemporary Finland where investment in reproduction is far lower, and modern medicine may alleviate many potential costs. Finally, we would like to acknowledge that due to unaccounted variation in individual heterogeneity, our study will still underestimate the underlying trade-off, and future research should focus on the underlying mechanisms or within-individual measures of aging (e.g., (61)). Lifespan is the ultimate outcome of many underlying factors, and reproductive behavior in particular can have different effects on different diseases: for example, high reproductive effort increases the risk of cardiovascular disease (47) but very low reproductive effort may increase the risk of breast cancer (62). Therefore, future studies focusing on specific diseases or using biomarkers would be best placed to study how individual reproductive behavior shapes aging.

## MATERIALS AND METHODS

### Study population

To investigate how a mother’s reproductive behavior affects her lifespan, we used long-term individual life-history data from rural Finland. Data were adapted from Parish records across rural Finland dating back to the 16^th^ century. These records were maintained by the Lutheran Church and include birth, marriage, and death certificates, but also yearly attendances of religious services, which from 1749 onwards were required by law to be recorded for tax purposes. This allows complete life-histories to be precisely constructed, including the migration of individuals between parishes. These individual records were then linked across generations resulting in a genealogical dataset of 100,598 people born from 1536 to 2012.

The majority of individuals in these data were born in a preindustrial period where agriculture employed 90% of the population (35). Preindustrial farming techniques, combined with particularly tough crop growing conditions in Finland, meant that during this period yearly yields of the two main crops, rye and barley, varied a lot with the climatic conditions (33). Consequently, the population was characterized by high birth and death rates due to limited access to healthcare and contraception, and infectious diseases such as smallpox, typhus, and typhoid fever (58). In our records, mothers on average had 4 children, but 38% of children died before their 15^th^ birthday. On average, women had their first child at age 25 and 95% had their last child before age 45. Like other pre-industrial European populations, the mating system was patrilocal and highly monogamous. Divorce was forbidden (63) and infidelity severely punished (64), with recent genetic studies showing that ∼1% of children were born outside of marital ties (65,66).

Industrialization began in Finland in the 1870s – relatively late by European standards – and developed gradually, with manufacturing still employing only 20% of the population by 1910 (35). Throughout this period, access to healthcare improved and vaccination of children against smallpox was relatively common by the 1880s (50). Consequently, life expectancy in Finland increased from 32 years at birth in 1865 to 70 for those born in 1965 and is now above the European average (67). In this period, access to contraception also improved and cultural preferences changed, with the fertility transition beginning around 1910 where birth rates declined from ∼30 births per 1000 people to ∼12 by the 1970s (49).

### Quantification of lifespans

All data handling was performed in R 4.3.2 (68) within RStudio 2023.12.1 (69), using *tidyr* 1.3.1 (70). We calculated the lifespan of all individuals with a recorded birth (99%) and death (43%). For those with recorded years but missing months and/or days the middle value was used (1% and 2% of recorded births and deaths, respectively). Individuals with missing death dates were then assigned a last-seen date of the final year they were known to have been alive in the data (available for 97% of individuals missing death dates), which could include a last-known birth of child or attendance of religious services. As the precise date of these services was often unknown, we standardized the date across individuals and treated all individuals as at least being alive on January 1^st^ of their year of last appearance. Lifespans were then calculated in decimal years using the *lubridate* package 1.9.2 (71) giving a median of 24 (range = 0-108). We also recorded whether this lifespan was calculated from known death dates or dates of last appearance which would be accounted for using a survival analysis (see *Statistical analyses*).

### Quantification of reproductive traits

We recorded six aspects of a woman’s reproductive history thought to capture a mother’s reproductive effort, defined as the total amount of resources devoted to bearing and raising offspring to independence (38). These variables included the four variables that Helle (38) used to measure reproductive effort: number of children born, number of child-years experienced, ages at last reproduction, and reproductive tenure (i.e., the time between first and last reproduction). Additionally, we included the number of sons born, and the number of children born that were twins or triplets (hence multiple births), which may be associated with increased reproductive effort (72). We also modified the definition of child-years experienced from the cumulative time that a mother co-survived with her children while they were younger than age 18 to age 5, because the effort of bearing and raising offspring is relatively low and children are more independent after age 5 (73), even helping to raise younger siblings thereafter (74). Thus, if a mother survived at least 5 years after her last birth and had 3 children, one of which died at age 1, another age 10, and another age 60, her child years experienced would be (1+5+5=) 11.

The median number of children for known mothers in our population was 3, ranging from 1 to 18. We estimated the ages at last reproduction and reproductive tenure for all mothers with a known child-birth date, which had medians of 35 (range, 16–51) and 8 (0–31), respectively. Sex of children was known in 99.4% of children, with mothers having 0–12 sons, with population-level birth sex ratios approximately 50:50. Multiple birth status was assigned to 99.3% of cases. Six percent of mothers had at least one set of twins or triplets, but one mother had 3 sets of twins. In cases where children or mothers had unknown death dates before age 5 (81%), the year of last appearance was used to calculate the child-years experienced. Median child years experienced were 12 (0-75).

### Assigning famine exposure groups

All individuals were grouped according to whether – and in what life stage – they were exposed to the Great Finnish Famine. We first assigned the start and end dates of the famine as February 1^st^ 1865 and May 31^st^ 1869 which is the period of elevated excess mortality observed country-wide in Finland due to the famine (35). Based on these dates we categorized all women surviving to adulthood (19.1yr, 5^th^ percentile for age at first reproduction) into 3 groups: individuals dying before the famine (n = 3,694), individuals exposed to the famine at some point in their live (n = 9,337), and individuals born after the famine (n = 15,036). We then further divided individuals exposed to the famine into three groups based on when they were exposed to the famine: when they were developing, reproductively active, or post-reproductive. We defined women as reproductively active during the famine if – at the midpoint of the famine (i.e., March 31^st^ 1867) – they were older than the 5^th^ percentile of age at first reproduction (19.1 yr) but younger than the 95^th^ percentile of age at last reproduction (44.9 yr) calculated from mothers reproducing before the fertility transition in 1911 (49). Individuals younger than 19.1 years at the famine’s midpoint were assigned the developing group (n = 4,231) and individuals older than 44.9 at the midpoint were assigned the post-reproductive group (n = 1,612), with those aged in between assigned to the reproductively active group (n = 3,494). 1,052 women could not be assigned a famine category, due to missing birth or last seen dates.

### Assigning socioeconomic status groups

To account for a source of individual heterogeneity that can influence the lifespan and reproductive effort of women (48), we grouped women surviving to adulthood into three socioeconomic status groups: upper, including mostly landowners and merchants (n = 6,591); intermediate, including mostly craftsmen, such as smiths, and fishermen (n = 7,764); and lower, consisting of landless families and servants (n = 6,750). In cases where a woman’s occupation was missing but the husband’s occupation was present, we assigned it based on the husband’s occupation (n = 2,357). If both were present but different (n = 590), the highest socioeconomic status was used. In total, socioeconomic status could be determined for 76% of adult women.

### Data selection

The data selection is summarized in Fig. S2. For women surviving to adulthood, we first excluded childless women (n = 9,275) because of the substantial differences in health of childless females in preindustrial populations that make comparisons to reproductive females (henceforth mothers) challenging (12). We then excluded mothers that had not been tracked throughout their entire reproductive period (n = 6,658) to ensure the precise estimation of reproductive effort. We further excluded mothers who died before completing their reproductively active period (age 44.9, as defined above) because we wanted to estimate the long-term lifespan costs of reproduction and minimize frailty heterogeneity (n = 2,204). However, we accept that this may lead to an underestimation of the total costs of reproduction by excluding those who died directly after reproducing and by including only the most robust mothers that survived to post-reproductive life (23,40). This left 9,930 women. Before 1725 the coverage of the data was very low and after 1911 there were increasing numbers of missing individuals because they are still alive, both of which could bias lifespans, so were removed (n = 1,906). We were also only interested in the long-term effects of the famine and so excluded those dying during the famine period itself (n = 478). We further controlled for individual heterogeneity by excluding mothers who were twins or triplets (n = 119), multiple marriers (n = 612), and women without recorded childbirth dates (n = 4) or socioeconomic statuses (n = 410) as done in other studies (75).

Finally, prior to analyses we examined the proportion of individuals with known death dates versus those that had only a last seen date (i.e. were censored) across the years lived for each of our famine exposure groups. Survival analyses (as used here) allow the combining of censored and uncensored data to estimate lifespans but can become biased if the proportion of censoring is biased with respect to lifespan across predictors in a survival model (76). We found that the percentage of censored individuals was much higher in the groups of individuals exposed to the famine while reproductively active (43%), while developing (51%), and those born after the famine (35%), compared with those exposed post-reproduction (4%) and dying before the famine (3%; Fig. S3a), due to two spikes in the number of censored individuals between the years of 1898-1904 and 1910-11 (Fig. S3b). Because of the temporal nature of our famine exposure groups, these resulted in systematic structuring of censoring across lifespans (Fig. S4), which would result in biased lifespans. Removing lifespans censored during those two periods (n = 1717) removed these biases (Fig. S4) and resulted in a relatively even percentage of censored to non-censored individuals across famines exposure categories for analyses (2-11%; Fig. S5).

This resulted in 4684 mothers with a median lifespan of at least 68 years (range 45-103) and 5 children (1-17) (Table S6). These values are slightly higher than the full dataset (see Table S6) but remained comparable to other similar populations (22,75).

### Statistical analysis

Following Helle (38) we used a confirmatory factor analysis measurement model to estimate the *latent* lifetime reproductive effort from the six indicator variables that are hypothesized to be components of a woman’s reproductive effort (77–79). Measurement models use simultaneous equations to separate the correlation between these indicator variables into the true variance in reproductive effort, and the error variance which captures the measurement error for each specific indicator variable (80). As indicator variables we used the number of children born, number of child-years experienced, ages at last reproduction, and reproductive tenure, number of sons born, and number of children that were part of multiple births. Indicators with factor loading p-values < 0.05 contributed to estimating lifetime reproductive effort. The number of children born was used to scale the *latent* reproductive effort so that each unit increase in reproductive effort could be interpreted as equivalent to the total resources used in raising one child. The latent reproductive effort was mean centered at 0 so values could be negative, and indicate lower than average reproductive effort.

All reproductive variables significantly contributed to estimating reproductive effort (p-value < 0.001, *loadings*, Table 1). Specifically, increases in one unit of reproductive effort was associated with 0.536 (95% CIs = 0.523-0.548) more sons, 1.238 (1.187-1.289) years increase in ages at last reproduction, 2.131 (2.092-2.170) years longer reproductive tenures, 3.878 (3.811-3.945) years increase in child-years experienced, and 0.052 (0.044-0.059) more multiple births (Table 1). The number of children was clearly the most important component of the reproductive effort variable and showed a correlation with the estimated reproductive effort of 0.99 (see Fig. S6).

This measurement model was incorporated within a structural equation model (SEM) estimated using simultaneous equations and restricted maximum likelihood in the software *Mplus* 8.8 (81). This model was used to examine how reproductive effort, lifespan, and the direct effect of reproductive effort on lifespan changed across mothers exposed to the Great Finnish Famine. Lifespan was estimated using a semiparametric continuous-time survival model which approximates a Cox-proportional hazards model (82). This combines the number of years individuals were known to have lived and whether these individuals had a recorded death date or had been censored. By using a survival analysis, the SEM estimates how the log-hazard mortality risk changes according to each covariate. Hence, positive coefficients indicate an increased risk of mortality with an increase in the covariate, and thus a reduction in survival probability or lifespan. The model included the direct effects of reproductive effort on lifespan while simultaneously estimating the differences in reproductive effort and lifespan across predictors. Separating these effects is a key strength of the SEM, which cannot be done in other models (such as linear mixed models). We also modelled the differences in lifespan and reproductive effort across socioeconomic statuses and temporal changes occurring in each famine exposure group by including a birth year effect relative to the middle birth year in each famine exposure group, and by allowing the effects of relative birth year to vary across groups. We also tested if the effect of reproductive effort on lifespan depended on socioeconomic status. P-values less than 0.05 were deemed significant and non-significant interactions were removed from final models, least significant first, to aid interpretation of lower-order effects. Pairwise comparisons between categories were done by swapping the reference groups in the SEM. Collinearity of between variables used in models was checked using *corvif ()* from *Highstat* 10 (83). The variance inflation factors were all < 1.5. For illustration and quantification purposes, we extracted the estimated reproductive effort for mothers and ran Cox proportional hazard models with the same predictors for each famine exposure group separately, allowing us to estimate and present the predicted effect of changing reproductive effort on lifespan under a given famine exposure.

*Mplus* does not allow for the inclusion of random effects if some groups only have one sample (81). Therefore, we also used the extracted reproductive effort estimates for each mother from the SEM in mixed effect Cox proportional hazard models using *Coxme* 2.2-20 (84) to see if our results were robust when accounting for shared family, region, and birth year effects among mothers. For these analyses we retained all but four individuals of unknown region. Mothers with unknown parents were assumed to have come from different families (44%) as sisters would have been identifiable in the data. *Coxme* models cannot model the direct effects of factors such as famine exposure on reproductive effort, but they represent a useful robustness check and still account for such dependencies when estimating the marginal effects. For *Coxme* models, we used likelihood ratio chi-squared tests comparing models with and without a given predictor to test for the significance of predictors with a threshold of p < 0.05.

The SEMs were run in *Mplus* (81), implemented in R 4.3.0 (68) using *MplusAutomation* 1.1.0 (85). *ggplot2* 3.3.5 (86) and *ggpubr* 0.4.0 (87) were used for plotting, *ggeffects* 1.5.1 (88) was used for estimating marginal effects. *survival* 3.5.7 (89) was used for estimating life-expectancies which were calculated as the predicted age at which survival probability was 50%, using median values for variables and an intermediate socioeconomic status if not otherwise specified.

## COMPETING INTERESTS

The authors declare no competing interests.

## FUNDING

E.A.Y.’s PhD was supported by the University of Groningen, through a Rosalind Franklin Fellowship awarded to H.L.D. V.L. was funded by the Strategic Research Council of the Academy of Finland (grant nos. 345185 and 345183).

## DATA AVAILABILITY STATEMENT

Code and data necessary for reproducing results are available at: https://doi.org/10.34894/HEWSRP

## AUTHOR CONTRIBUTION

Conceptualization and methodology: EAY, EP, VL, HLD

Data collection: VL

Data wrangling and analysis: EAY

Writing—original draft: EAY

Writing—review & editing: EAY, EP, VL, HLD

## ACKNOWLEDGEMENTS

We thank Mirkka Lahdenpera, Mila Salonen, Dinos Sevdalakis, Aida Nitsch, and the University of Exeter’s Quantitative Genetics Club for feedback throughout the project. We thank Samuli Helle for helping with the SEM approach and other detailed methodological feedback, and Yuheng Sun, Maaike A Versteegh, Heung Ying Janet Chik, Marianthi Tangili, and Barbara Tschirren for comments on the writing.

## Notes

### Competing Interest Statement

The authors have declared no competing interest.

### Summary of Updates

Accepted version. Some changes have been made to the text, mainly the introduction and discussion.

## REFERENCES

1. Flatt T, Partridge L. Horizons in the evolution of aging. BMC Biol [Internet]. 2018 Dec [cited 2024 Nov 12];16(1):93. Available from: https://bmcbiol.biomedcentral.com/articles/10.1186/s12915-018-0562-z

2. Kirkwood TBL. Evolution of ageing. Nature. 1977;270(5635):301–4. Available from: 10.1038/270301a0

3. Kirkwood TBL, Rose MR. Evolution of senescence: late survival sacrificed for reproduction. Philosophical Transactions: Biological Sciences. 1991;332(1262):15–24. Available from: 10.1098/rstb.1991.0028

4. Flatt T, Heyland A, editors. Mechanisms of Life History Evolution: The Genetics and Physiology of Life History Traits and Trade-Offs [Internet]. Oxford University Press; 2011 [cited 2024 Nov 12]. Available from: https://academic.oup.com/book/10632

5. Chang C, Moiron M, Sánchez-Tójar A, Niemelä PT, Laskowski KL. What is the meta-analytic evidence for life-history trade-offs at the genetic level? Ecology Letters [Internet]. 2024 Jan [cited 2024 Nov 12];27(1):e14354. Available from: https://onlinelibrary.wiley.com/doi/10.1111/ele.14354

6. Winder LA, Simons MJ, Burke T. No evidence for a trade-off between reproduction and survival in a meta-analysis across birds. eLife [Internet]. 2025 Mar 31 [cited 2025 Apr 21];12:RP87018. Available from: https://elifesciences.org/articles/87018

7. Boonekamp JJ, Salomons M, Bouwhuis S, Dijkstra C, Verhulst S. Reproductive effort accelerates actuarial senescence in wild birds: an experimental study. Sorci G, editor. Ecology Letters [Internet]. 2014 May [cited 2024 Sept 3];17(5):599–605. Available from: https://onlinelibrary.wiley.com/doi/10.1111/ele.12263

8. Jasienska G. Costs of reproduction and ageing in the human female: Reproduction and ageing in women. Philosophical Transactions of the Royal Society B: Biological Sciences. 2020;375(1811). Available from: https://royalsocietypublishing.org/doi/10.1098/rstb.2019.0615

9. Beeton M, Yule U, Pearson K. Data for the problem of evolution in man. V. On the correlation between duration of life and the number of offspring. Proc R Soc Lond [Internet]. 1901 Feb 28 [cited 2024 Sept 3];67(435–441):159–79. Available from: https://royalsocietypublishing.org/doi/10.1098/rspl.1900.0015

10. Bolund E. The challenge of measuring trade-offs in human life history research. Evolution and Human Behavior [Internet]. 2020;41(6):502–12. Available from: 10.1016/j.evolhumbehav.2020.09.003

11. Gagnon A. Natural fertility and longevity. Fertility and Sterility [Internet]. 2015;103(5):1109–16. Available from: 10.1016/j.fertnstert.2015.03.030

12. Hurt L, Ronsmans C, Thomas SL. The effect of number of births on women’s mortality: Systematic review of the evidence for women who have completed their childbearing. Population Studies [Internet]. 2006;60(1):55–71. Available from: https://www.semanticscholar.org/paper/5f7ddbc9e722b95a3fb093545d99697640c03637

13. Cohen AA, Coste CFD, Li XY, Bourg S, Pavard S. Are trade-offs really the key drivers of ageing and life span? Functional Ecology. 2020;34(1):153–66. Available from: 10.1111/1365-2435.13444

14. Reznick D, Nunney L, Tessier A. Big houses, big cars, superfleas and the costs of reproduction. Trends in Ecology and Evolution [Internet]. 2000;15(10):421–5. Available from: 10.1016/S0169-5347(00)01941-8

15. van Noordwijk AJ, de Jong G. Acquisition and Allocation of Resources: Their Influence on Variation in Life History Tactics. The American Naturalist [Internet]. 1986;128(1):137–42. Available from: 10.1086/284547

16. Sear R. The impact of reproduction on Gambian women: Does controlling for phenotypic quality reveal costs of reproduction? American Journal of Physical Anthropology. 2007;132(4):632–41.

17. Doblhammer G, Oeppen J. Reproduction and longevity among the British peerage: The effect of frailty and health selection. Proceedings of the Royal Society B: Biological Sciences [Internet]. 2003;270(1524):1541–7. Available from: 10.1098/rspb.2003.2400

18. Messina FJ, Fry J. Environment-dependent reversal of a life history trade-off in the seed beetle Callosobruchus maculatus. Journal of evolutionary biology [Internet]. 2003;16(3):501–9. Available from: 10.1046/j.1420-9101.2003.00535.x

19. Bliard L, Martin JS, Paniw M, Blumstein DT, Martin JGA, Pemberton JM, Nussey DH, Childs DZ, Ozgul, A. Detecting context dependence in the expression of life history trade-offs. Journal of Animal Ecology [Internet]. 2025 Mar [cited 2025 Aug 26];94(3):379–93. Available from: 10.1111/1365-2656.14173

20. Dribe M. Long-term effects of childbearing on mortality: Evidence from pre-industrial Sweden. Popul. Population Studies [Internet]. 2004;58(3):297–310. Available from: 10.1080/0032472042000272357

21. Lycett JE, Dunbar RIM, Voland E. Longevity and the costs of reproduction in a historical human population. Proceedings of the Royal Society B: Biological Sciences [Internet]. 2000;267(1438):31–5. Available from: 10.1098/rspb.2000.0962

22. Invernizzi L, Bergeron P, Pelletier F, Lemaître JF, Douhard M. Sons Shorten Mother’s Lifespan in Preindustrial Families with a High Level of Infant Mortality. The American Naturalist [Internet]. 2024 Aug 21 [cited 2024 Sept 2];204(4):315–26. Available from: https://www.journals.uchicago.edu/doi/10.1086/731792

23. Hsu CH, Posegga O, Fischbach K, Engelhardt H. Examining the trade-offs between human fertility and longevity over three centuries using crowdsourced genealogy data. PLoS ONE [Internet]. 2021;16(8 August):1–20. Available from: 10.1371/journal.pone.0255528

24. Nenko I, Hayward AD, Simons MJP, Lummaa V. Early-life environment and differences in costs of reproduction in a preindustrial human population. PLoS ONE. 2018;13(12):1–16. Available from: 10.1371/journal.pone.0207236

25. Wang X, Byars S, Stearns S. Genetic links between post-reproductive lifespan and family size in Framingham. Evolution Medicine & Public Health [Internet]. 2013;2013(1):241–53. Available from: https://www.semanticscholar.org/paper/91a5850557983601dccf100dfd78e1da8e11f1f9

26. Riley JC. Rising life expectancy: a global history. Cambridge, UK: Cambridge University Press; 2001.

27. Vermeulen R, Schymanski EL, Barabási AL, Miller GW. The exposome and health: Where chemistry meets biology. Science [Internet]. 2020 Jan 24 [cited 2024 Nov 21];367(6476):392–6. Available from: https://www.science.org/doi/10.1126/science.aay3164

28. Cheng M, Conley D, Kuipers T, Li C, Ryan CP, Taeubert MJ, Wang S, Wang T, Zhou J, Schmitz J, Schmitz LL, Tobi EW, Heijmans BT, Lumey LH, Belsky, DW. Accelerated biological aging six decades after prenatal famine exposure. Proceedings of the National Academy of Sciences. 2024;121(24):e2319179121. Available from: 10.1073/pnas.2319179121

29. Lumey LH, Li C, Khalangot M, Levchuk N, Wolowyna O. Fetal exposure to the Ukraine famine of 1932–1933 and adult type 2 diabetes mellitus. Science [Internet]. 2024 Aug 8;385(6709):667–71. Available from: https://www.science.org/doi/10.1126/science.adn4614

30. Lumey LH, Stein AD, Susser E. Prenatal Famine and Adult Health. Annu Rev Public Health [Internet]. 2011 Apr 21 [cited 2023 May 9];32(1):237–62. Available from: https://www.annualreviews.org/doi/10.1146/annurev-publhealth-031210-101230

31. Chen Y, Zhou LA. The long-term health and economic consequences of the 1959–1961 famine in China. Journal of Health Economics [Internet]. 2007 July [cited 2024 Dec 22];26(4):659–81. Available from: 10.1016/j.jhealeco.2006.12.006

32. Barker DJP. Mothers, babies and disease in later life. Edinburgh; New York: Churchill Livingstone; 1998. 217 p.

33. Hayward AD, Holopainen J, Pettay JE, Lummaa V. Food and fitness: Associations between crop yields and life-history traits in a longitudinally monitored pre-industrial human population. Proceedings of the Royal Society B: Biological Sciences [Internet]. 2012;279(1745):4165–73. Available from: 10.1098/rspb.2012.1190

34. Rickard IJ, Holopainen J, Helama S, Helle S, Russell AF, Lummaa V. Food availability at birth limited reproductive success in historical humans. Ecology. 2010;91(12):3515–25. Available from: 10.1890/10-0019.1

35. Voutilainen M. Poverty, Inequality and the Finnish 1860s Famine. 2016. Available from: https://urn.fi/URN:ISBN:978-951-39-6627-0

36. Pitkänen KJ, Mielke JH. Age and sex differentials in mortality during two nineteenth century population crises. European Journal of Population. 1993;9(1):1–32. Available from: https://link.springer.com/10.1007/BF01267899

37. Salonen M, Lahdenperä M, Rotkirch A, Lummaa V. Fertility resilience varies by socioeconomic status and sex: Historical trends in childlessness across 150 years. iScience [Internet]. 2024 July 19;27(110227):1–13. Available from: 10.1016/j.isci.2024.110227

38. Helle S. Search for a resource-based trade-off between lifetime reproductive effort and women’s postreproductive survival in preindustrial Sweden. Journals of Gerontology - Series A Biological Sciences and Medical Sciences [Internet]. 2019;74(5):642–7. Available from: 10.1093/gerona/gly203

39. Helle S, Lummaa V, Jokela J. Are reproductive and somatic senescence coupled in humans? Late, but not early, reproduction correlated with longevity in historical Sami women. Proceedings of the Royal Society B: Biological Sciences [Internet]. 2005;272(1558):29–37. Available from: 10.1098/rspb.2004.2944

40. Helle S. Selection bias in studies of human reproduction-longevity trade-offs. Proceedings of the Royal Society B: Biological Sciences. 2017;284(1868). Available from: 10.1098/rspb.2017.2104

41. Jasienska G. Reproduction and lifespan: Tradeoffs, overall energy budgets, intergenerational costs, and costs neglected by research. American Journal of Human Biology [Internet]. 2009;21(4):524–32. Available from: 10.1002/ajhb.20931

42. Butte NF, King JC. Energy requirements during pregnancy and lactation. Public Health Nutr [Internet]. 2005 Oct [cited 2024 Sept 25];8(7a):1010–27. Available from: https://www.cambridge.org/core/product/identifier/S136898000500131X/type/journal_article

43. Jasienska G. Reproduction and lifespan: Trade-offs, overall energy budgets, intergenerational costs, and costs neglected by research. Am J Hum Biol [Internet]. 2009 July [cited 2023 May 25];21(4):524–32. Available from: https://onlinelibrary.wiley.com/doi/10.1002/ajhb.20931

44. Saltiki K, Alevizaki M. Coronary heart disease in postmenopausal women; the role of endogenous estrogens and their receptors. Hormones [Internet]. 2007;6(1):9–24. Available from: http://www.hormones.gr/pdf/Coronary%20heart%20disease.pdf

45. Heinig MJ, Dewey KG. Health effects of breast feeding for mothers: a critical review. Nutr Res Rev [Internet]. 1997 Jan [cited 2024 Oct 1];10(1):35–56. Available from: https://www.cambridge.org/core/product/identifier/S095442249700005X/type/journal_article

46. Shin YA, Lee KY. Low estrogen levels and obesity are associated with shorter telomere lengths in pre- and postmenopausal women. J Exerc Rehabil [Internet]. 2016 June 30 [cited 2024 Oct 1];12(3):238–46. Available from: http://e-jer.org/journal/view.php?number=2013600273

47. Li W, Ruan W, Lu Z, Wang D. Parity and risk of maternal cardiovascular disease: A dose– response meta-analysis of cohort studies. Eur J Prev Cardiolog [Internet]. 2019 Apr [cited 2023 Aug 11];26(6):592–602. Available from: https://academic.oup.com/eurjpc/article/26/6/592-602/5925653

48. Pettay JE, Helle S, Jokela J, Lummaa V. Natural Selection on Female Life-History Traits in Relation to Socio-Economic Class in Pre-Industrial Human Populations. PLoS ONE [Internet]. 2007;2(7). Available from: 10.1371/journal.pone.0000606

49. Mitchell B. International Historical Statistics, Brian Mitchell (2013) – processed by Our World in Data. “International Historical Statistics (Births per 1,000) (Brian Mitchell (2013))” [dataset]. International Historical Statistics, Brian Mitchell (2013) [original data]. 2013.

50. Ukonaho S, Lummaa V, Briga M. The Long-Term Success of Mandatory Vaccination Laws After Implementing the First Vaccination Campaign in 19th Century Rural Finland. American Journal of Epidemiology [Internet]. 2022 June 27 [cited 2023 Dec 4];191(7):1180–9. Available from: https://academic.oup.com/aje/article/191/7/1180/6549054

51. Kaptijn R, Thomese F, Liefbroer AC, Van Poppel F, Van Bodegom D, Westendorp RGJ. The Trade-Off between Female Fertility and Longevity during the Epidemiological Transition in the Netherlands. Sear R, editor. PLoS ONE [Internet]. 2015 Dec 17 [cited 2023 May 22];10(12):e0144353. Available from: https://dx.plos.org/10.1371/journal.pone.0144353

52. Doblhammer G, Van Den Berg GJ, Lumey LH. A re-analysis of the long-term effects on life expectancy of the Great Finnish Famine of 1866–68. Population Studies [Internet]. 2013 Nov 1 [cited 2023 May 2];67(3):309–22. Available from: https://www.tandfonline.com/doi/full/10.1080/00324728.2013.809140

53. Hayward AD, Rickard IJ, Lummaa V. Influence of early-life nutrition on mortality and reproductive success during a subsequent famine in a preindustrial population. Proceedings of the National Academy of Sciences of the United States of America [Internet]. 2013;110(34):13886–91. Available from: 10.1073/pnas.1301817110

54. Hammel EA, Gullickson A. Kinship structures and survival: Maternal mortality on the Croatian–Bosnian border 1750–1898. Population Studies [Internet]. 2004 July [cited 2024 Oct 1];58(2):145–59. Available from: http://www.tandfonline.com/doi/abs/10.1080/0032472042000213703

55. Saxton K, Falconi A, Goldman-Mellor S, Catalano R. No evidence of programmed late-life mortality in the Finnish famine cohort. J Dev Orig Health Dis [Internet]. 2013 Feb [cited 2024 Aug 27];4(1):30–4. Available from: https://www.cambridge.org/core/product/identifier/S2040174412000517/type/journal_article

56. Hughes K, Bellis MA, Hardcastle KA, Sethi D, Butchart A, Mikton C, Jones L, Dunne MP. The effect of multiple adverse childhood experiences on health: a systematic review and meta-analysis. The Lancet Public Health [Internet]. 2017 Aug [cited 2024 Nov 12];2(8):e356–66. Available from: https://linkinghub.elsevier.com/retrieve/pii/S2468266717301184

57. Cartwright SJ, Nicoll MAC, Jones CG, Tatayah V, Norris K. Anthropogenic Natal Environmental Effects on Life Histories in a Wild Bird Population. Current Biology [Internet]. 2014 Mar [cited 2024 Sept 24];24(5):536–40. Available from: https://linkinghub.elsevier.com/retrieve/pii/S0960982214000736

58. Turpeinen O. Regional differentials in finnish mortality rates 1816-1865. Scandinavian Economic History Review [Internet]. 1973;21(2):145–63. Available from: 10.1080/03585522.1973.10407768

59. Lummaa V. Early developmental conditions and reproductive success in humans: Downstream effects of prenatal famine birthweight, and timing of birth. American Journal of Human Biology. 2003;15(3):370–9. Available from: 10.1073/pnas.1301817110

60. Painter RC, Westendorp RGJ, De Rooij SR, Osmond C, Barker DJP, Roseboom TJ. Increased reproductive success of women after prenatal undernutrition. Human Reproduction [Internet]. 2008 July 29 [cited 2025 Sept 10];23(11):2591–5. Available from: https://academic.oup.com/humrep/article-lookup/doi/10.1093/humrep/den274

61. Ryan CP, Lee NR, Carba DB, MacIsaac JL, Lin DTS, Atashzay P, Belsky DW, Kober MS, Kuzawa CW. Pregnancy is linked to faster epigenetic aging in young women. Proc Natl Acad Sci USA [Internet]. 2024 Apr 16 [cited 2024 Sept 25];121(16):e2317290121. Available from: https://pnas.org/doi/10.1073/pnas.2317290121

62. Aktipis CA, Ellis BJ, Nishimura KK, Hiatt RA. Modern reproductive patterns associated with estrogen receptor positive but not negative breast cancer susceptibility. Evolution, Medicine and Public Health [Internet]. 2015;2015(1):52–74. Available from: 10.1093/emph/eou028

63. Moring B. Nordic family patterns and the north-west European household system. Continuity and Change [Internet]. 2003;18(1):77–109. Available from: 10.1017/S0268416003004508

64. Sundin J. Sinful Sex: Legal Prosecution of Extramarital Sex in Preindustrial Sweden. Social Science History [Internet]. 1992;16(1):99–128. Available from: 10.2307/1171323

65. Larmuseau MHD, Matthijs K, Wenseleers T. Cuckolded Fathers Rare in Human Populations. Trends in Ecology and Evolution [Internet]. 2016;31(5):327–9. Available from: 10.1016/j.tree.2016.03.004

66. Larmuseau MHD, van den Berg P, Claerhout S, Calafell F, Boattini A, Gruyters L, Vandenbosch M, Nivelle K, Decorte R, Wenseleers T. A Historical-Genetic Reconstruction of Human Extra-Pair Paternity. Current Biology [Internet]. 2019;29(23):4102–4107.e7. Available from: 10.1016/j.cub.2019.09.075

67. Wilmoth JR, Andreev K, Jdanov D, Glei DA, Riffe T, Boe C, Bubenheim M, Philipov D, Shkolnikov V, Vachon P, Winant C, Barberi M. Human Mortality Database. Available at www.mortality.org.; 2021.

68. R Core Team. R: A language environment for statistical computing [Internet]. Vienna, Austria: R Foundation for Statistical Computing; 2020. Available from: http://www.r-project.org/.

69. Allaire J. RStudio: integrated development environment for R. Boston, MA. 2012;770(394):165–71.

70. Wickham H, Averick M, Bryan J, Chang W, McGowan LD, François R, Grolemund G, Hayes A, Henry L, Hester J, Kuhn M, Pedersen, TL, Miller E, Bache SM, Müller K, Ooms J, Robinson, David, Seidel DP, Spinu V, Takahashi K, Vaughan D, Wilke C, Woo K, Yutani H. Welcome to the tidyverse. Journal of Open Source Software [Internet]. 2019;4(43):1686. Available from: https://joss.theoj.org/papers/10.21105/joss.01686

71. Grolemund G, Wickham H. Dates and Times Made Easy with {lubridate}. Journal of Statistical Software [Internet]. 2011;40(3):1–25. Available from: https://www.jstatsoft.org/v40/i03/

72. Invernizzi L, Lemaître J, Douhard M. The expensive son hypothesis. Journal of Animal Ecology [Internet]. 2024 Oct 30 [cited 2024 Dec 2];1365-2656.14207. Available from: https://besjournals.onlinelibrary.wiley.com/doi/10.1111/1365-2656.14207

73. Riswick T, Hsieh YH. Between rivalry and support: The impact of sibling composition on infant and child mortality in Taiwan, 1906‒1945. DemRes [Internet]. 2020 Mar 24 [cited 2023 Nov 3];42:615–56. Available from: https://www.demographic-research.org/volumes/vol42/21/

74. Crognier E, Villena M, Vargas E. Helping patterns and reproductive success in Aymara communities. Am J Hum Biol [Internet]. 2002 May [cited 2023 Nov 9];14(3):372–9. Available from: https://onlinelibrary.wiley.com/doi/10.1002/ajhb.10047

75. Gagnon A, Smith KR, Tremblay M, Vézina H, Paré PP, Desjardins B. Is there a trade-off between fertility and longevity? A comparative study of women from three large historical databases accounting for mortality selection. Am J Hum Biol [Internet]. 2009 July [cited 2023 May 16];21(4):533–40. Available from: https://onlinelibrary.wiley.com/doi/10.1002/ajhb.20893

76. Leung KM, Elashoff RM, Afifi AA. Censoring issues in survival analyses. Annu Rev Public Health [Internet]. 1997 May [cited 2023 Aug 29];18(1):83–104. Available from: https://www.annualreviews.org/doi/10.1146/annurev.publhealth.18.1.83

77. Bollen K. Structural Equations With Latent Variables [Internet]. New York: Wiley; 1989. Available from: http://doi/book/10.1002/9781118619179

78. Heck RH, Thomas SL. Chapter 5 Defining multilevel latent variables. In: An introduction to multilevel modelling techniques: MLM and SEM approaches using Mplus [Internet]. 3rd edn New York: Routledge; 2015. p. 133–81. Available from: 10.4324/9781315746494

79. Muthén BO. Beyond SEM: General Latent Variable Modeling. Behaviormetrika [Internet]. 2002 Jan [cited 2024 Jan 17];29(1):81–117. Available from: http://link.springer.com/10.2333/bhmk.29.81

80. Kilne R. Principles and Practice of Structural Equation Modeling [Internet]. 4th edn. New York: The Guilford Press; 2016. Available from: https://www.statmodel.com/download/SurvivalJSM3.pdf

81. Muthén LK, Muthén BO. Mplus: User’s Guide [Internet]. Los Angeles, CA: Muthén & Muthén; 1998. 201 p. Available from: https://www.mendeley.com/library/

82. Asparouhov T, Muth B. Continuous - Time Survival Analysis in Mplus. 2018;(1):1–22.

83. Zuur AF, Ieno EN, Walker NJ, Saveliev AA, Smith GM, others. Mixed effects models and extensions in ecology with R [Internet]. Vol. 574. Springer; 2009. Available from: https://link.springer.com/book/10.1007/978-0-387-87458-6

84. Therneau TM. coxme: Mixed Effects Cox Models [Internet]. 2024. Available from: https://CRAN.R-project.org/package=coxme

85. Hallquist MH, Wiley JF. MplusAutomation: An R Package for Facilitating Large-Scale Latent Variable Analyses in Mplus. Structural Equation Modeling: A Multidisciplinary Journal [Internet]. 2018;25(4):621–38. Available from: 10.1080/10705511.2017.1402334

86. Wickham H. ggplot2: Elegant Graphics for Data Analysis. 2016; Available from: https://ggplot2.tidyverse.org

87. Kassambara Alboukadel. ggpubr: ‘ggplot2’ Based Publication Ready Plots [Internet]. 2020. Available from: https://cran.r-project.org/package=ggpubr

88. Lüdecke D. ggeffects: Tidy Data Frames of Marginal Effects from Regression Models. JOSS [Internet]. 2018 June 29 [cited 2024 Jan 19];3(26):772. Available from: http://joss.theoj.org/papers/10.21105/joss.00772

89. Therneau T. A Package for Survival Analysis in R. R package version 3.2-7. 2020. URL https://CRANR-project.org/package=survival. 2020;

